# Protein kinases mediate anti-inflammatory effects of cannabidiol and estradiol against high glucose in cardiac sodium channels

**DOI:** 10.1101/2020.11.23.395129

**Authors:** Mohamed A. Fouda, Peter C. Ruben

## Abstract

**Background and purpose:** Cardiovascular anomalies are predisposing factors for diabetes-induced morbidity and mortality. Recently, we showed that high glucose induces changes in the biophysical properties of Nav1.5 that could be strongly correlated to diabetes-induced arrhythmia. However, the mechanisms underlying hyperglycemia-induced inflammation, and how inflammation provokes cardiac arrhythmia, are not well understood. We hypothesized that inflammation could mediate the high glucose-induced biophyscial changes on Nav1.5 through protein phosphorylation by protein kinases A and C. We also hypothesized that this signaling pathway is, at least partly, involved in the cardiprotective effects of CBD and E_2_.

**Experimental approach:** To test these ideas, we used Chinese hamster ovarian (CHO) cells transiently co-transfected with cDNA encoding human Nav1.5 α-subunit under control, a cocktail of inflammatory mediators or 100 mM glucose conditions (for 24 hours). We used electrophysiological experiments and action potential modelling.

**Key Results:** Inflammatory mediators, similar to 100 mM glucose, right shifted the voltage dependence of conductance and steady state fast inactivation and increased persistent current leading to computational prolongation of action potential (hyperexcitability) which could result in long QT3 arrhythmia. In addition, activators of PK-A or PK-C replicated the inflammation-induced gating changes of Nav1.5. Inhibitors of PK-A or PK-C, CBD or E_2_ mitigated all the potentially deleterious effects provoked by high glucose/inflammation.

**Conclusions and implications:** These findings suggest that PK-A and PK-C may mediate the anti-inflammatory effects of CBD and E_2_ against high glucose-induced arrhythmia. CBD, via Nav1.5, may be a cardioprotective therapeutic approach in diabetic postmenopausal population.

**Bullet points:** **What is already known:**

- Arrhythmias are among the common cardiac causes of morbidity and mortality in diabetes-related hyperglycemia.
- One of the diabetes-induced arrhythmias is long-QT syndrome, caused by gating defects in the cardiac voltage-gated sodium channel (Nav1.5).

**What this study adds:**

- Inflammation and subsequent activation of PK-A and PK-C mediate the high glucose-induced electrophysiological changes of Nav1.5 in a manner consistent with the gating defects that underlie long-QT arrhythmia.
- Cannabidiol and estradiol rescue the high glucose induced Nav1.5 gating defects through, at least partly, this signaling pathway.

**Clinical significance:**

- Inflammation/PK-A and PK-C signaling pathway could be a potential therapeutic target to prevent arrhythmias associated with diabetes.
- Cannabidiol may be a therapeutic approach to prevent cardiac complications in diabetes, especially in postmenopausal populations due to the decreased levels of the cardioprotective estrogen.

## Introduction

Cardiovascular anomalies are strongly correlated with diabetes-induced morbidity and mortality (Matheus, Tannus, Cobas, Palma, Negrato & Gomes, 2013). These deleterious cardiovascular complications are mainly attributed to hyperglycemia/high glucose (Pistrosch, Natali & Hanefeld, 2011). There is also a positive correlation between diabetes/high glucose and long QT (LQT) syndrome (Fouda, Ghovanloo & Ruben, 2020; Grisanti, 2018). LQT syndrome is a cardiac arrhythmogenic disorder, identified by a prolongation of the Q-T interval. One cause of LQT syndrome is a gain-of-function in cardiac sodium channels, as in LQT3 (Shimizu & Antzelevitch, 1999).

Oxidative stress and activation of pro-inflammatory pathways are among the main pathways involved in diabetes/high glucose evoked cardiovascular abnormalities (Rajesh et al., 2010). Cardiac inflammation has a key role in the development of cardiovascular anomalies (Adamo, Rocha-Resende, Prabhu & Mann, 2020). Inhibition of inflammatory signaling pathways ameliorate cardiac consequences (Adamo, Rocha-Resende, Prabhu & Mann, 2020). Ion channels are crucial players in inflammation-induced cardiac abnormalities (Eisenhut & Wallace, 2011). Voltage-gated sodium channels (Nav) underlie phase 0 of the cardiac action potential (Balser, 1999; Ruan, Liu & Priori, 2009). Changes in the biophysical properties of the primary cardiac sodium channel, Nav1.5, are linked to diabetes induced cardiovascular abnormalities (Fouda, Ghovanloo & Ruben, 2020; Yu et al., 2018). However, the mechanisms underlying hyperglycemia-induced inflammation, and how inflammation provokes cardiac dysfunction, are not well understood.

Cannabidiol (CBD) is approved as an anti-seizure drug (Barnes, 2006; Devinsky et al., 2017). CBD lacks adverse cardiac toxicity and ameliorates diabetes/high glucose induced deletrious cardiomyopathy (Cunha et al., 1980; Izzo, Borrelli, Capasso, Di Marzo & Mechoulam, 2009; Rajesh et al., 2010). Recently, we showed that CBD rescues the biophysical substrate for LQT3 via direct inhibitory effects on cardiac sodium ion channels and indirect anti-oxidant effects (Fouda, Ghovanloo & Ruben, 2020). In addition, CBD inhibits the production of pro-inflammatory cytokines *in vitro* and *in vivo* (Nichols & Kaplan, 2020).

Gonadal hormones have crucial roles in the inflammatory responses (El-Lakany, Fouda, El-Gowelli, El-Gowilly & El-Mas, 2018; El-Lakany, Fouda, El-Gowelli & El-Mas, 2020). Estrogen (E_2_), the main female sex hormone, acts via genomic and non-enomic mechanisms to inhibit inflammatory cascades (Murphy, Guyre & Pioli, 2010). Clinically, postmenopausal females exhibited higher levels of TNF-α in reponse to endotoxemia compared with pre-menopausal women (Moxley, Stern, Carlson, Estrada, Han & Benson, 2004). Interestingly, E_2_ stabilizes Nav fast inactivation and reduces the late sodium currents (Wang, Garro & Kuehl-Kovarik, 2010), similar to CBD effects on Nav1.5 (Fouda, Ghovanloo & Ruben, 2020).

Here, we characterized the role of inflammation in high glucose-induced biophyscial changes on Nav1.5. Second, we found that changes in the biophysical properties of Nav1.5 may be, at least in part, mediated through protein phosphorylation by protein kinases A and C. Finally, we show that this signaling pathway may be, at least partly, involved in the cardiprotective effects of CBD and E_2_.

## Materials and Methods

### Cell culture

Chinese hamster ovary cells (CHO) (RRID: CVCL_0214) were grown at pH 7.4 in filtered sterile F12 (Ham’s) nutrient medium (Life Technologies, Thermo Fisher Scientific, Waltham, MA, USA), supplemented with 5% FBS and maintained in a humidified environment at 37°C with 5% CO2. Cells were transiently co-transfected with the human cDNA encoding the Nav1.5 α-subunit, the β1-subunit, and eGFP. Transfection was done according to the PolyFect (Qiagen, Germantown, MD, USA) transfection protocol. A minimum of 8-hour incubation was allowed after each set of transfections. The cells were subsequently dissociated with 0.25% trypsin– EDTA (Life Technologies, Thermo Fisher Scientific) and plated on sterile coverslips under normal (10 mM) or elevated glucose concentrations (100 mM) (Fouda, Ghovanloo & Ruben, 2020) or a cocktail of inflammatory mediators (Akin et al., 2019) containing bradykinin (1 μM), PGE-2 (10 μM), histamine (10 μM), 5-HT (10 μM), and adenosine 5’-triphosphate (15 μM) for 24 hours prior to electrophysiological experiments.

### Electrophysiology

Whole-cell patch clamp recordings were made using an extracellular solution composed of NaCl (140 mM), KCl (4 mM), CaCl_2_ (2 mM), MgCl_2_ (1 mM), HEPES (10 mM). The extracellular solution was titrated to pH 7.4 with CsOH. Pipettes were fabricated with a P-1000 puller using borosilicate glass (Sutter Instruments, CA, USA), dipped in dental wax to reduce capacitance, then thermally polished to a resistance of 1.0–1.5 MΩ. Pipettes were filled with intracellular solution, containing: CsF (120 mM), CsCl (20 mM), NaCl (10 mM), HEPES (10 mM) titrated to pH 7.4. All recordings were made using an EPC-9 patch-clamp amplifier (HEKA Elektronik, Lambrecht, Germany) digitized at 20 kHz via an ITC-16 interface (Instrutech, Great Neck, NY, USA). Voltage clamping and data acquisition were controlled using PatchMaster/FitMaster software (HEKA Elektronik, Lambrecht, Germany) running on an Apple iMac (Cupertino, California). Current was low-pass-filtered at 5 kHz. Leak subtraction was automatically done using a P/4 procedure following the test pulse. Gigaohm seals were allowed to stabilize in the on-cell configuration for 1 min prior to establishing the whole-cell configuration. Series resistance was less than 5 MΩ for all recordings. Series resistance compensation up to 80% was used when necessary. All data were acquired at least 5 min after attaining the whole-cell configuration, and cells were allowed to incubate 5 min after drug application prior to data collection. Before each protocol, the membrane potential was hyperpolarized to −130 mV to insure complete removal of both fast-inactivation and slow-inactivation. Leakage and capacitive currents were subtracted with a P/4 protocol. All experiments were conducted at 22 °C.

### Activation protocols

To determine the voltage-dependence of activation, we measured the peak current amplitude at test pulse voltages ranging from −130 to +80 mV in increments of 10 mV for 19 ms. Channel conductance (G) was calculated from peak I_Na_:

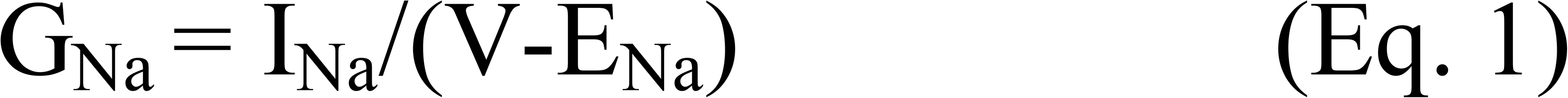

where G_Na_ is conductance, I_Na_ is peak sodium current in response to the command potential V, and E_Na_ is the Nernst equilibrium potential. The midpoint and apparent valence of activation were derived by plotting normalized conductance as a function of test potential. Data were then fitted with a Boltzmann function:

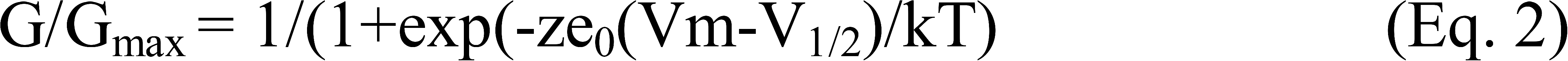

where G/G_max_ is normalized conductance amplitude, Vm is the command potential, z is the apparent valence, e_0_ is the elementary charge, V_1/2_ is the midpoint voltage, k is the Boltzmann constant, and T is temperature in K.

### Steady state fast inactivation protocols

The voltage-dependence of fast-inactivation was measured by preconditioning the channels to a hyperpolarizing potential of −130 mV and then eliciting pre-pulse potentials that ranged from −170 to +10 mV in increments of 10 mV for 500 ms, followed by a 10 ms test pulse during which the voltage was stepped to 0 mV. Normalized current amplitude as a function of voltage was fit using the Boltzmann function:

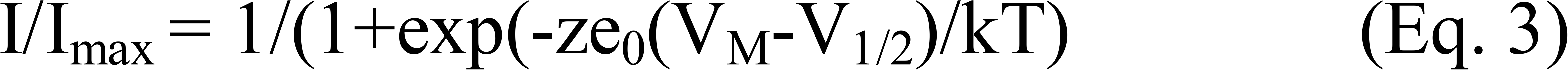

where I_max_ is the maximum test pulse current amplitude. z is apparent valency, e_0_ is the elementary charge, Vm is the prepulse potential, V_1/2_ is the midpoint voltage of SSFI, k is the Boltzmann constant, and T is temperature in K.

### Fast inactivation recovery

Channels were fast inactivated during a 500 ms depolarizing step to 0 mV. Recovery was measured during a 19 ms test pulse to 0 mV following −130 mV recovery pulse for durations between 0 and 1.024 s. Time constants of fast inactivation were derived using a double exponential equation:

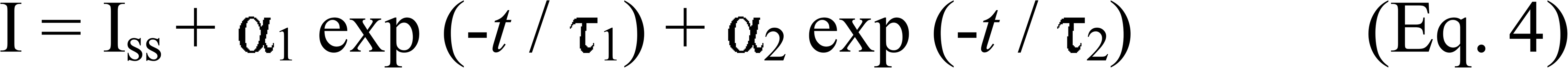

where I is current amplitude, I_ss_ is the plateau amplitude, α_1_ and α_2_ are the amplitudes at time 0 for time constants τ_1_ and τ_2_, and *t* is time.

### Persistent current protocols

Late sodium current was measured between 45 and 50◻ms during a 50◻ms depolarizing pulse to 0◻mV from a holding potential of −130◻mV. Fifty pulses were averaged to increase signal to noise ratio (Abdelsayed, Peters & Ruben, 2015; Abdelsayed, Ruprai & Ruben, 2018).

### Action potential modeling

Action potentials were simulated using a modified version of the O’Hara-Rudy model programmmed in Matlab (O’Hara et al. 2011, PLoS Comput. Bio). The code that was used to produce model is available online from the Rudy Lab website (http://rudylab.wustl.edu/research/cell/code/Allcodes.html). The modified gating INa parameters were in accordance with the biophysical data obtained from whole-cell patch-clamp experiments in this study for various conditions. The model accounted for activation voltage-dependence, steady-state fast inactivation voltage-dependence, persistent sodium currents, and peak sodium currents (compound conditions).

### Drug preparations

CBD was purchased from Toronto Research Chemicals (Toronto, Ontario) in powder form. Other compounds (e.g. 17β-estradiol (E_2_), bradykinin, PGE-2, histamine, 5-HT, adenosine 5’-triphosphate, D-glucose, Gö 6983 (PKC inhibitor), H-89 (PKA inhibitor), 8-(4-chlorophenylthio) adenosine-3’,5’-cyclic monophosphate (CPT-cAMP; PKA activator) or PMA (PKC activator)) were purchased from Sigma-Aldrich (ON, Canada). Powdered CBD, Gö 6983, H-89, adenosine CPT-cAMP or PMA were dissolved in 100% DMSO to create stock. The stock was used to prepare drug solutions in extracellular solutions at various concentrations with no more than 0.5% total DMSO content.

### Data analysis and statistics

The data and statistical analysis comply with the British Journal of Pharmacology recommendations on experimental design and analysis in pharmacology (Curtis et al., 2018). Studies were designed to generate groups of almost equal size (n=5), using randomisation and blinded analysis. Normalization was performed in order to control the variations in sodium channel expression and inward current amplitude and in order to be able to fit the recorded data with Boltzmann function (for voltage-dependences) or an exponential function (for time courses of inactivation). Fitting and graphing were done using FitMaster software (HEKA Elektronik, Lambrecht, Germany) and Igor Pro (Wavemetrics, Lake Oswego, OR). Statistical analysis consisted of one-way ANOVA (endpoint data) along with post hoc testing of significant findings along with Student’s t-test and Tukey’s test using Prism 7 software (Graphpad Software Inc., San Diego, CA). Values are presented as mean ± SEM with probability levels less than 0.05 considered significant. Statistical analysis was undertaken only for studies where each group size was at least “n=5”. The declared group size is the number of independent values, and that statistical analysis was done using these independent values. In the electrophysiological experiments, we randomized the different treatments under the different conditions (e.g. control vs. high glucose or inflammatory mediators), so that five cells in each treatment or condition came from five different randomized cell passages.

## Results

### Inflammatory mediators alter the gating properties of Nav1.5 similar to high glucose

We recently showed that high glucose, in a concentration-dependent manner, right shifts the voltage dependence of activation and steady state fast inactivation and increases persistent current (Fouda, Ghovanloo & Ruben, 2020). Here, we used whole-cell voltage-clamp to measure gating in human Nav1.5, and test the effects of incubating for 24 hours in either a cocktail of inflammatory mediators (Akin et al., 2019) or 100 mM glucose (Fouda, Ghovanloo & Ruben, 2020). Peak channel conductance was measured between −130 and +80 mV. We measured channel conductance in the presence of inflammatory mediators to determine whether the high glucose induced-changes in Nav1.5 activation (Fouda, Ghovanloo & Ruben, 2020) are, at least partly, mediated through inflammation. Fig. 1A shows conductance plotted as a function of membrane potential. High glucose (100 mM) significantly shifted the Nav1.5 midpoint (V_1/2_) of activation in the positive direction (P= 0.0002). Additionally, the slope (apparent valence, z) of the activation curves showed a significant decrease in 100 mM glucose (P=0.007) (Fig. 1A and Table 1). This decrease in slope suggests a reduction in activation charge sensitivity. We found that incubation in inflammatory mediators for 24 hours, similar to 100 mM glucose, significantly right-shifted V_1/2_ of activation (P=0.001) and decreased z of activation curve (P=0.03) (Fig. 1A and Table 1). This suggests that both 100 mM glucose or inflammatory mediators decrease the probability of Nav1.5 activation.

**Figure 1.**
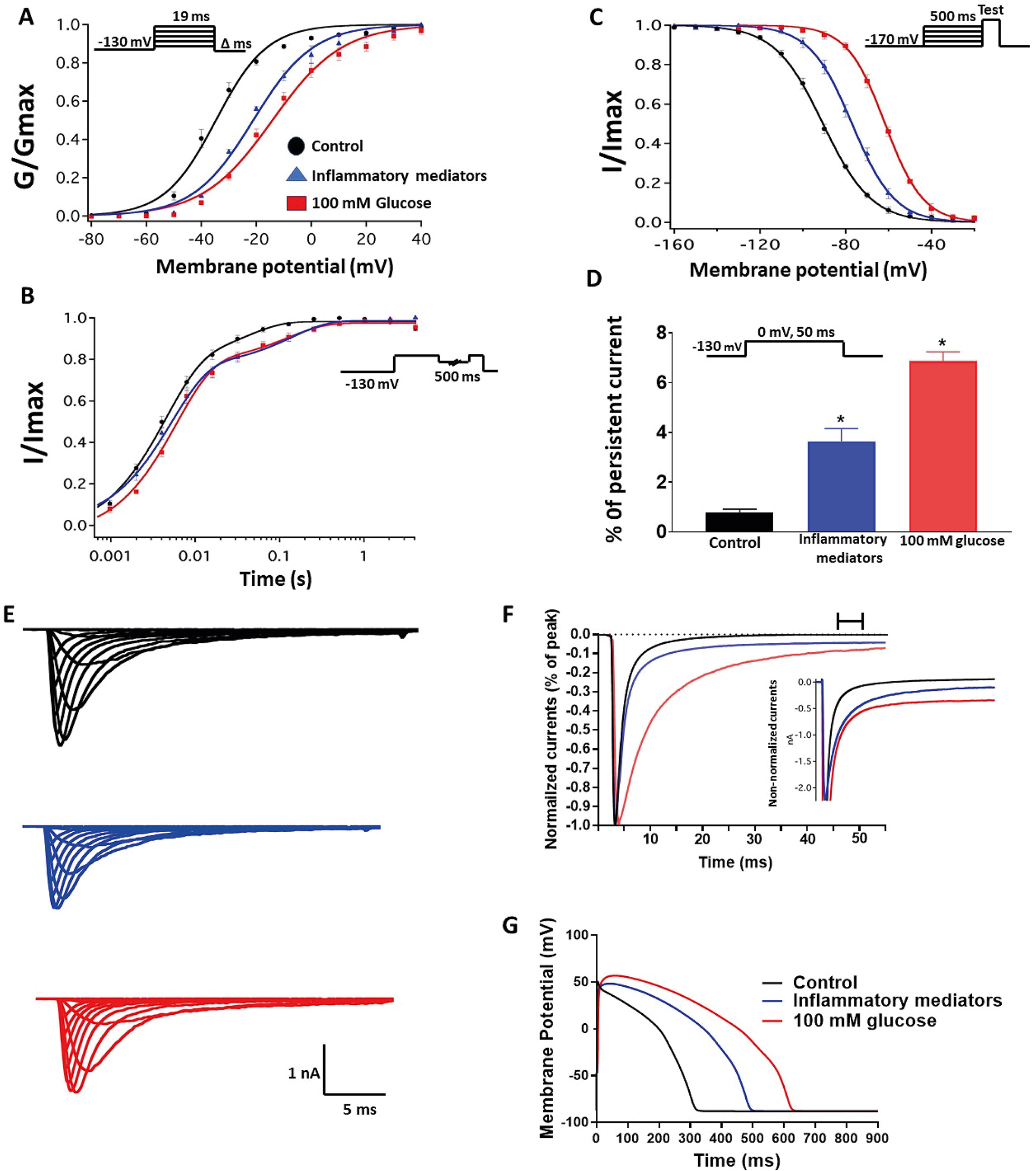
(A) Effect of a cocktail of inflammatory mediators or 100 mM glucose or their vehicle (for 24 hours) on conductance curve of Nav1.5 transfected cells with the insert showing the protocol (n=5, each). (B) Effect of a cocktail of inflammatory mediators or 100 mM glucose or their vehicle (for 24 hours) on SSFI of Nav1.5 transfected cells with the insert showing the protocol (n=5, each). (C) Effect of a cocktail of inflammatory mediators or 100 mM glucose or their vehicle (for 24 hours) on recovery from fast inactivation of Nav1.5 transfected cells with the insert showing the protocol (n=5, each). (D) Effect of a cocktail of inflammatory mediators or 100 mM glucose or their vehicle (for 24 hours) on the percentage of persistent sodium currents of Nav1.5 transfected cells with the insert showing the protocol (n=5, each). (E) Representative families of macroscopic currents. (F) Representative persistent currents across conditions. Currents were normalized to peak current amplitude. Bar above current traces indicates period during which persistent current was measured. Inset shows non-normalized currents. (G) *In silico* action potential duration of Nav1.5 transfected cells incubated in inflammatory mediators or 100 mM glucose or the vehicle for 24 hours. \**P* < 0.05 versus corresponding “Control” values.

**Table 1:**
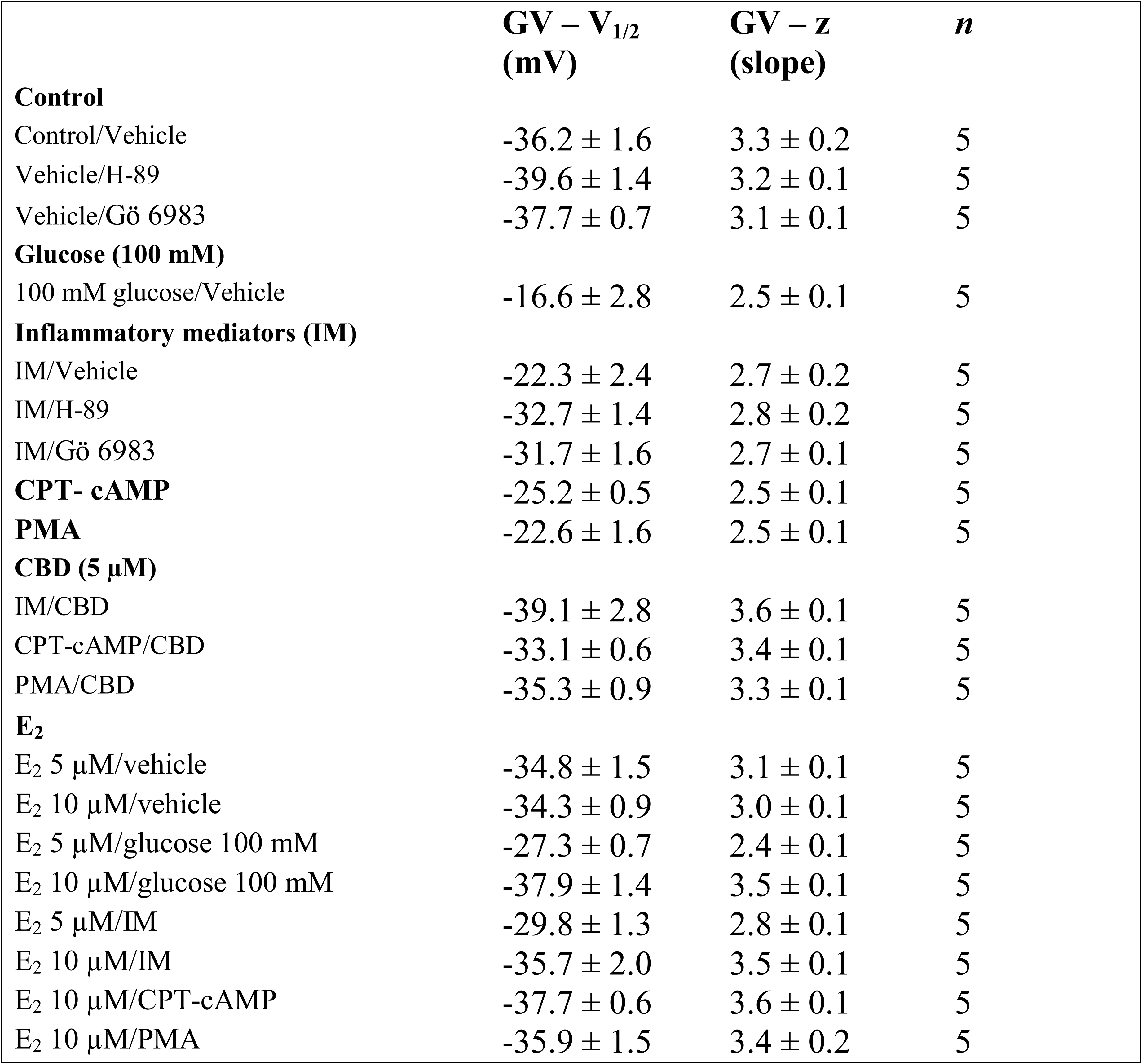
Steady state activation.

The DIII-IV linker mediates fast inactivation within a few milliseconds of Nav activation (West, Patton, Scheuer, Wang, Goldin & Catterall, 1992). Figure 1B shows normalized current amplitudes plotted as a function of pre-pulse potential. 100 mM glucose or inflammatory mediators caused significant shifts in the positive direction in the V_1/2_ obtained from Boltzmann fits (100 mM glucose: P<0.0001; inflammatory mediators: P=0.001) (Fig. 1B and Table 2). These shifts indicate a loss-of-function in fast inactivation and suggest that high glucose or inflammatory mediators decrease the probability of steady-state fast inactivation in Nav1.5.

**Table 2:**
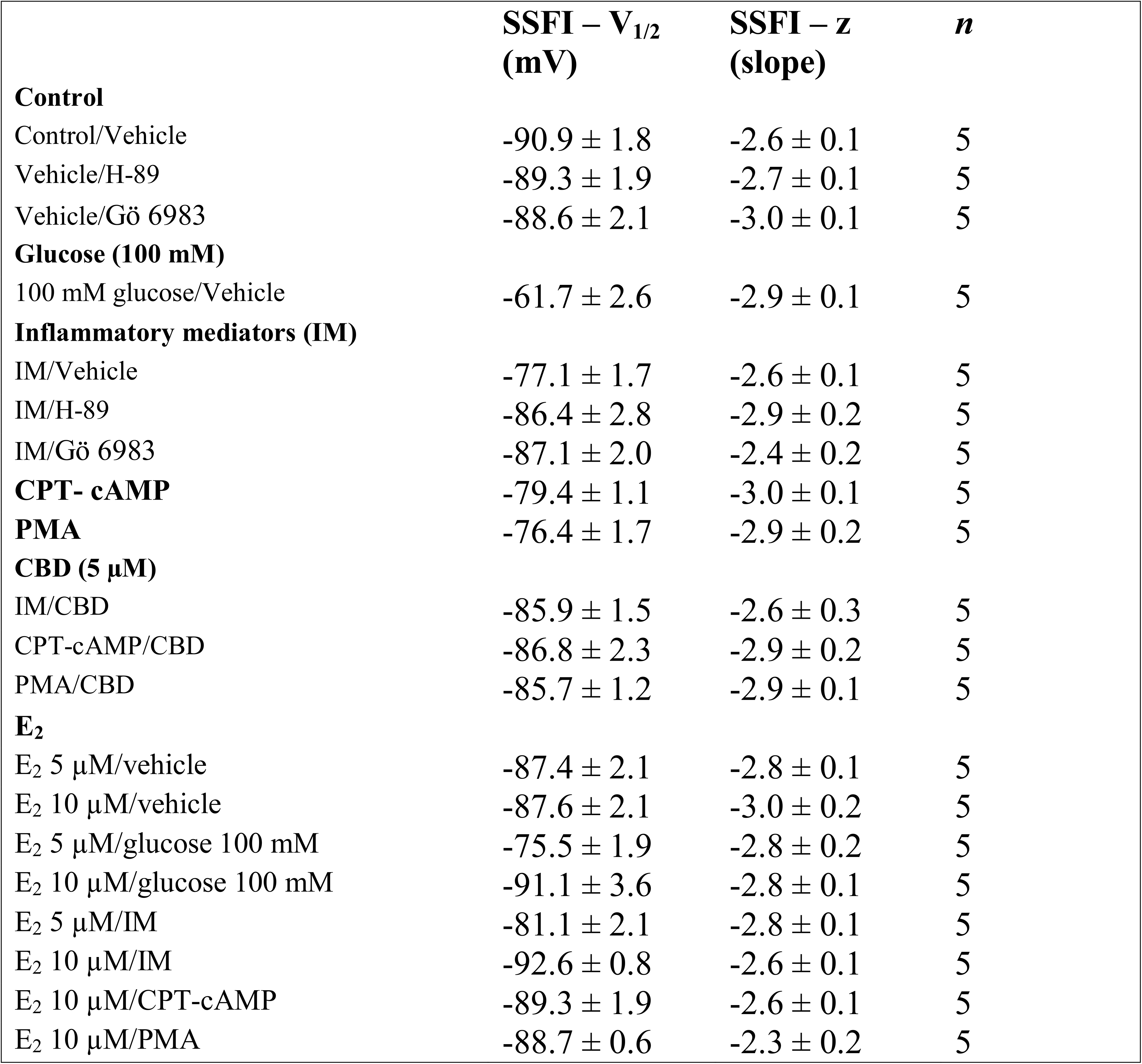
Steady state fast inactivation.

To measure fast inactivation recovery, we held channels at −130 mV to ensure channels were fully at rest, then depolarized the channels to 0 mV for 500 ms, and allowed different time intervals at −130 mV to measure recovery as a function of time. We found that incubation in 100 mM glucose or inflammatory mediators significantly (P<0.05) increase the slow component of fast inactivation recovery when compared to control, without affecting the fast component of recovery (Fig. 1C and Table 3).

**Table 3:**
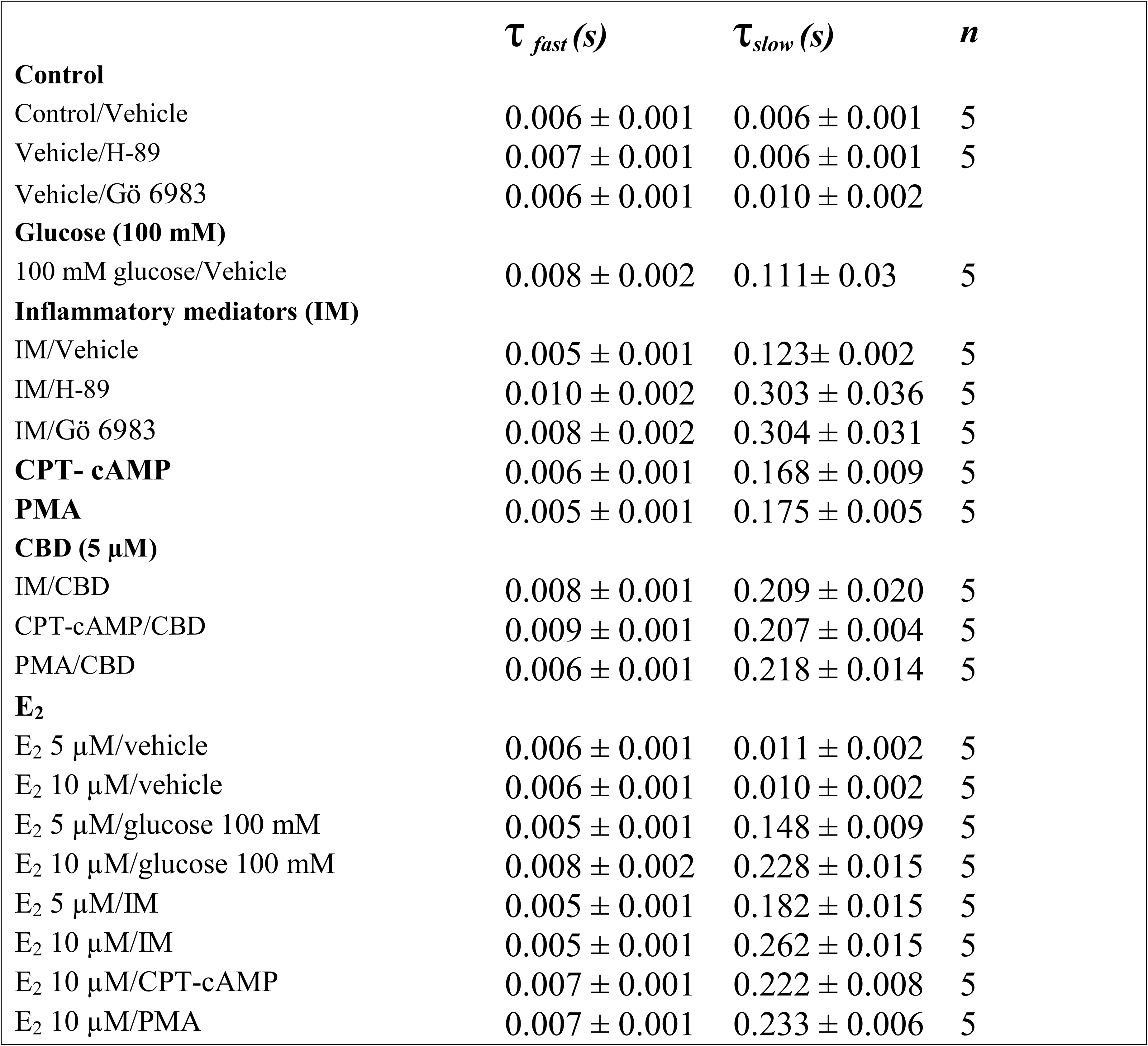
Time constants for the recovery from fast inactivation.

An increased persistent sodium current (INap) is a manifestation of destabilized fast inactivation (Goldin, 2003). Large INap is associated with a range of pathological conditions, including LQT3 (Ghovanloo, Abdelsayed & Ruben, 2016; Wang et al., 1995). To determine the effects of glucose or inflammatory mediators on the stability of Nav1.5 inactivation, we held channels at −130 mV, followed by a depolarizing pulse to 0 mV for 50 ms (Abdelsayed, Peters & Ruben, 2015; Abdelsayed, Ruprai & Ruben, 2018). Figure 1D shows that incubation in 100 mM glucose or inflammatory mediators significantly increased INap compared to control (100 mM glucose: P<0.0001; inflammatory mediators: P<0.0001) (Table 4). Representative families of macroscopic and persistent currents across conditions are shown (Fig. 1E and 1F).

**Table 4:**
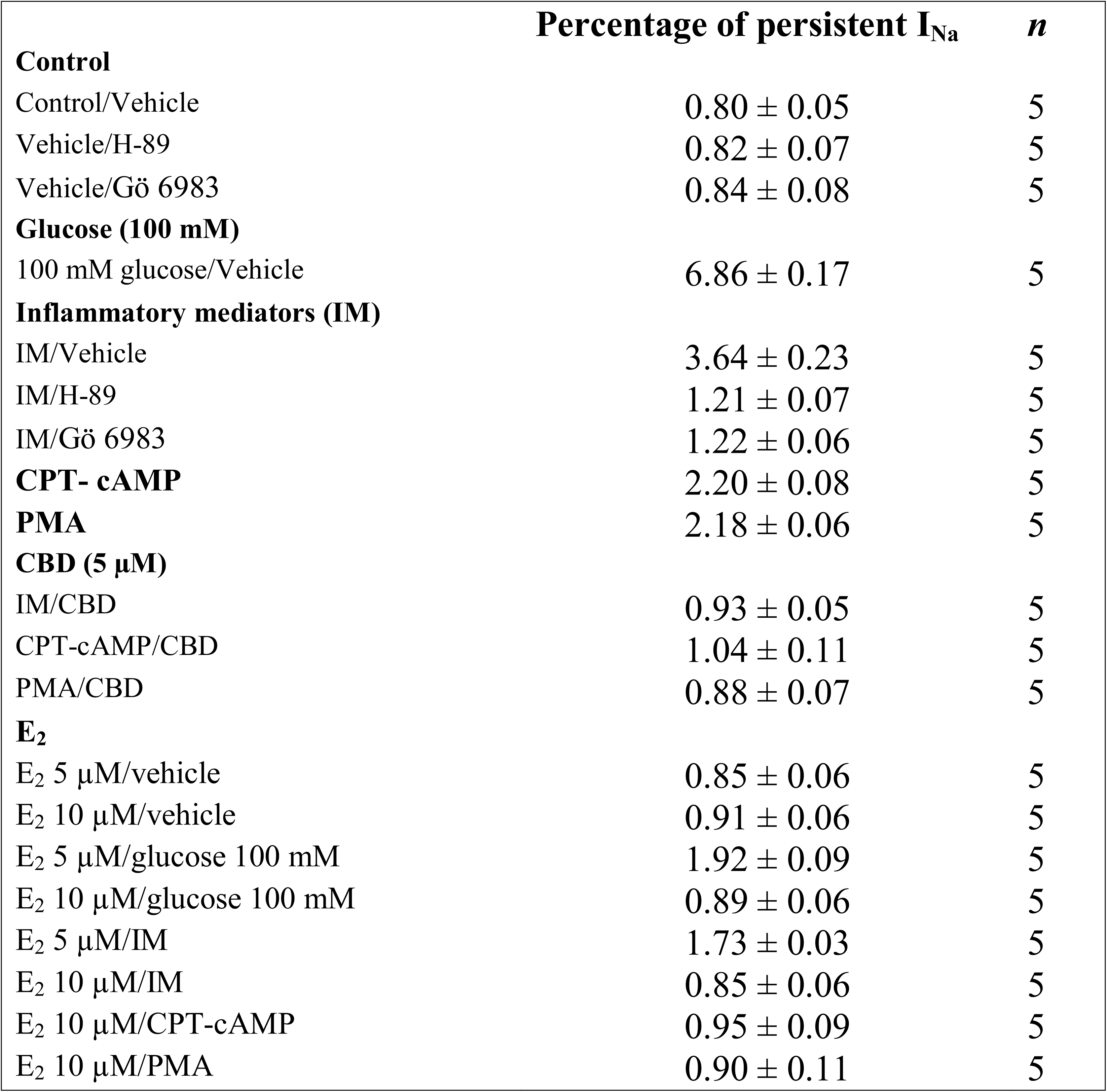
Persist ent current.

We used the O’Hara-Rudy model to simulate cardiac action potentials (AP) (O’Hara, Virag, Varro & Rudy, 2011). The model was modified using the results of our experiments and the effects of the tested compounds on the measured biophysical properties of activation (midpoint and apparent valence), steady-state fast inactivation (midpoint), recovery from fast inactivation, and persistent sodium current amplitude. The the original model parameters were adjusted to correspond to the control results from the patch-clamp experiments, and the subsequent magnitude shifts in the simulations of other conditions were performed relative to the control parameters (Fouda, Ghovanloo & Ruben, 2020). Figure 1G shows that modifying the model with data obtained from incubation in 100 mM glucose or inflammatory mediators prolonged the simulated AP duration (APD) from ~300 ms to ~500 ms (inflammatory mediators) and to > 600 ms (100 mM glucose). This increased APD potentially leads to the prolongation of the QT interval (Nachimuthu, Assar & Schussler, 2012). Despite the similarity between 100 mM glucose and inflammatory mediators-induced changes on Nav1.5, their responses are not exactly the same (Fig.1). This could be attributed to the concentration-dependent effects of high glucose on electrophysiological properties of Nav1.5 (Fouda, Ghovanloo & Ruben, 2020). We selected 100 mM glucose concentration to ensure a sufficiently large window to detect readout signals throughout the study.

### Activation of PK-A and PK-C mediates the inflammatory mediators induced alteration in the gating properties of Nav1.5

One of the key signaling pathways involved in inflammation is the activation of protein kinase A (PK-A) or protein kinases C (PK-C) and subsequent protein phosphorylation (Karin, 2005). To pharmacologically investigate the role of PK-A or PK-C signaling pathways in the inflammation-evoked gating changes of Nav1.5, we recorded Nav1.5 currents at room temperature in the absence, or after a 20 minute perfusion, of a PK-C activator (PMA; 10 nM (Hallaq, Wang, Kunic, George, Wells & Murray, 2012)) or PK-A activator (CPT-cAMP; 1 μM (Gu, Kwong & Lee, 2003; Ono, Fozzard & Hanck, 1993)). PMA or CPT-cAMP significantly shifted the Nav1.5 V_1/2_ of activation in the positive direction (PMA: P=0.0003; CPT-Camp: P=0.0007) (Fig 2A and table 1). In addition, PMA or CPT-Camp significantly reduced the effective valence (z) of the activation curves (PMA: P=0.002; CPT-Camp: P=0.007) (Fig. 2A and Table 1). Furthermore, PMA or CPT-cAMP caused significant right-shifts in the V_1/2_ of SSFI (PMA: P=0.0008; CPT-Camp: P=0.0005) (Fig. 2B and Table 2). Also, PMA or CPT-cAMP significantly (P<0.05) increase the slow component of fast inactivation recovery when compared to control (Fig. 2C and Table 3). We also found that PMA or CPT-cAMP significantly (PMA: P < 0.0001; CPT-Camp: P < 0.0001) increased INap compared to control (Fig. 2D and Table 4). These effects are similar to those of glucose and inflammatory mediators (Fig. 1). Representative families of macroscopic and persistent currents across conditions are shown (Fig. 2E and 2F). Similar to 100 mM glucose and inflammatory mediators, the data from PK-A (CPT-cAMP) or PK-C (PMA) activator experiments shows that the *in silico* APD increased from ~300 ms to ~400 ms (Fig. 2G).

**Figure 2.**
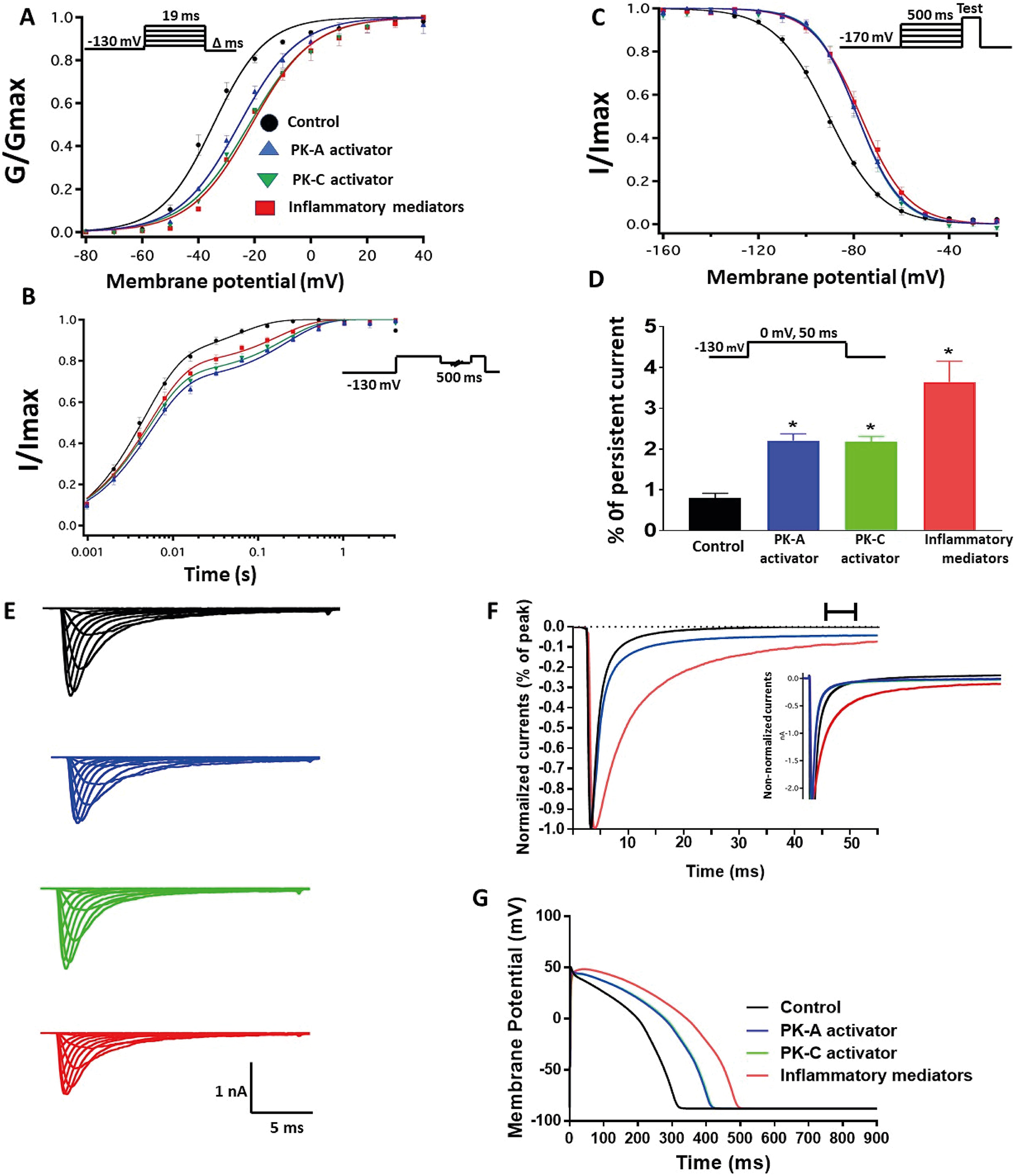
(A) Effect of inflammatory mediators (for 24 hours) or PK-A activator (CPT-cAMP; 1 μM, for 20 minutes) or PK-C activator (PMA; 10 nM, for 20 minutes) on conductance curve Nav1.5 transfected cells with the insert showing the protocol (n=5, each). (B) Effect of inflammatory mediators (for 24 hours) or PK-A activator (CPT-cAMP; 1 μM, for 20 minutes) or PK-C activator (PMA; 10 nM, for 20 minutes) on SSFI of Nav1.5 transfected cells with the insert showing the protocol (n=5, each). (C) Effect of inflammatory mediators (for 24 hours) or PK-A activator (CPT-cAMP; 1 μM, for 20 minutes) or PK-C activator (PMA; 10 nM, for 20 minutes) on recovery from fast inactivation of Nav1.5 transfected cells with the insert showing the protocol (n=5, each). (D) Effect of inflammatory mediators (for 24 hours) or PK-A activator (CPT-cAMP; 1 μM, for 20 minutes) or PK-C activator (PMA; 10 nM, for 20 minutes) on the percentage of persistent sodium currents of Nav1.5 transfected cells with the insert showing the protocol (n=5, each). (E) Representative families of macroscopic currents. (F) Representative persistent currents across conditions. Currents were normalized to peak current amplitude. Bar above current traces indicates period during which persistent current was measured. Inset shows non-normalized currents. (G) Effect of PK-A activator (CPT-cAMP; 1 μM for 20 minutes), PK-C activator (PMA; 10 nM, for 20 minutes) or inflammatory mediators (for 24 hours) on the *In silico* action potential duration of Nav1.5 transfected cells. \**P* < 0.05 versus corresponding “Control” values.

To ensure that the effects of the inflammatory mediators on Nav1.5 are indeed mediated, at least partly, through activation of PK-A and/or PK-C, we examined the effect of perfusing PK-A inhibitor (H-89, 2 μM for 20 minutes (Wang et al., 2013)) or PK-C inhibitor (Gö 6983, 1 μM for 20 minutes (Wang et al., 2013)) on Nav1.5 that had been incubated for 24 hours in either inflammatory mediators or vehicle. Although H-89 or Gö 6983 had no significant effects on Nav1.5 gating under control conditions (Table 1–4), H-89 or Gö 6983 reduced the inflammatory mediator-induced shifts in V_1/2_ (H-89: P=0.0108; Gö 6983: P=0.0203) (Fig. 3A and Table 1). In addition, H-89 or Gö 6983 rescued the inflammatory mediator-induced shift in Nav1.5 SSFI (Fig. 3B and Table 2). Moreover, H-89 or Gö 6983 (P=0.0041, or P=0.0017, respectively) further increased the time constant of the slow component of recovery from fast inactivation when compared to inflammatory mediators (Fig. 3C and Table 3).

**Figure 3.**
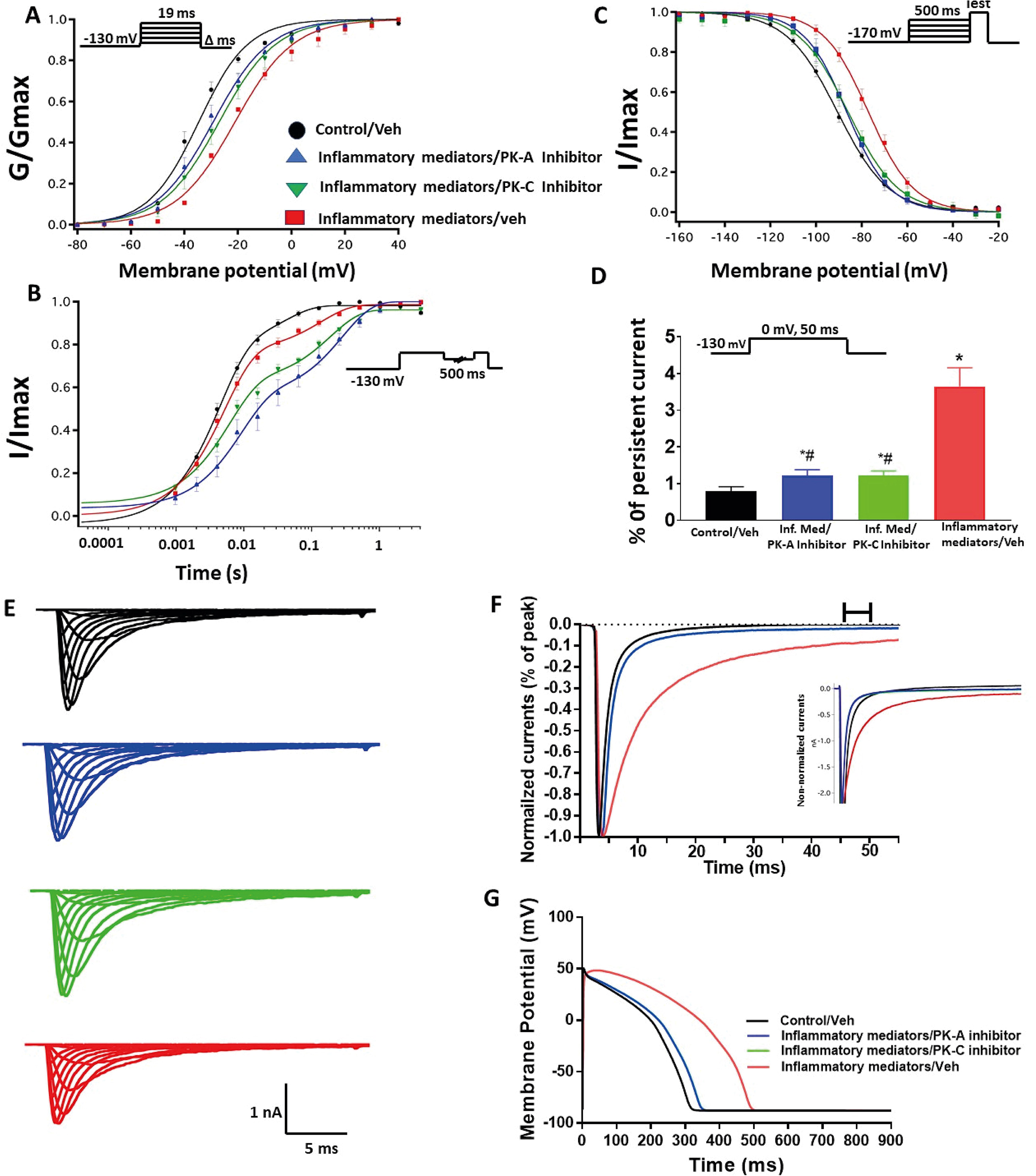
(A) Effect of PK-A inhibitor (H-89, 2 μM for 20 minutes) or PK-C inhibitor (Gö 6983, 1 μM for 20 minutes) or their vehicle on the conductance curve Nav1.5 transfected cells incubated in the inflammatory mediators for 24 hours with the insert showing the protocol (n=5, each). (B) Effect of PK-A inhibitor (H-89, 2 μM for 20 minutes) or PK-C inhibitor (Gö 6983, 1 μM for 20 minutes) or their vehicle on SSFI of Nav1.5 transfected cells incubated in the inflammatory mediators for 24 hours with the insert showing the protocol (n=5, each). (C) Effect of PK-A inhibitor (H-89, 2 μM for 20 minutes) or PK-C inhibitor (Gö 6983, 1 μM for 20 minutes) or their vehicle on recovery from fast inactivation of Nav1.5 transfected cells incubated in the inflammatory mediators for 24 hours with the insert showing the protocol (n=5, each). (D) Effect of PK-A inhibitor (H-89, 2 μM for 20 minutes) or PK-C inhibitor (Gö 6983, 1 μM for 20 minutes) or their vehicle on the percentage of persistent sodium currents of Nav1.5 transfected cells incubated in the inflammatory mediators for 24 hours with the insert showing the protocol (n=5, each). (E) Representative families of macroscopic currents. (F) Representative persistent currents across conditions. Currents were normalized to peak current amplitude. Bar above current traces indicates period during which persistent current was measured Inset shows non-normalized currents. (G) Effect of PK-A inhibitor (H-89, 2 μM for 20 minutes) or PK-C inhibitor (Gö 6983, 1 μM for 20 minutes) on the *In silico* action potential duration of Nav1.5 transfected cells incubated in inflammatory mediators for 24 hours. \**P* < 0.05 versus corresponding “Control/Veh” values. ^#^*P* < 0.05 versus corresponding “inflammatory mediators/Veh” values.

Figure 3D shows that H-89 or Gö 6983 (P < 0.0001) incompeletly reduced the inflammatory mediator-induced increase in the persistent currents (Table 4). Representative families of macroscopic and persistent currents across conditions are shown (Fig. 3E and 3F). Importantly, *in silico* APD using the data from inhibitors of PK-A (H-89) or PK-C (Gö 6983) reduced the inflammatory mediators-induced simulated APD prolongation (Fig. 3G).

### CBD rescues the Nav1.5 gating changes of inflammatory mediators, activation of PK-A or PK-C

Coupled with our previous observation that CBD rescues high glucose-induced dysfunciton in Nav1.5 (Fouda et al., 2020), our results from the above experiments with PK-A or PK-C modulators prompted us to test the effects of CBD on the biophysical properties of Nav1.5 in the presence of inflammatory mediators, PK-C activator (PMA), or PK-A activator (CPT-cAMP). To determine whether the observed changes to activation and SSFI imparted by inflammatory mediators or activation of PK-A or PK-C could be rescued, we measured peak sodium currents in the presence of CBD. CBD (5 μM) perfusion abolished the effects of inflammatory mediators, PMA, or CPT-cAMP, including shifts of V_1/2_ of activation, z of activation, and the V_1/2_ of SSFI (Fig. 4A and 4B and Table 1 and 2). In addition, CBD significantly increased the time constant of the slow component of recovery from fast inactivation regardless of the concurrent treatment (inflammatory mediators, PMA or CPT-cAMP) (Figure 4C and Table 3). Also, CBD reduced the increase in INap caused by inflammatory mediators, PMA or CPT-cAMP (Figure 4D and Table 4, with representative macroscopic and persistent currents shown in Fig. 4E and 4F). The O’Hara-Rudy model results also suggest that CBD rescues the prolonged *in silico* APD caused by inflammatory mediators or activators of PK-A or PK-C to nearly that of the control condition (Fig. 4G). The reduction in APD is consistent with the anti-excitatory effects of CBD (Ghovanloo, Shuart, Mezeyova, Dean, Ruben & Goodchild, 2018).

**Figure 4.**
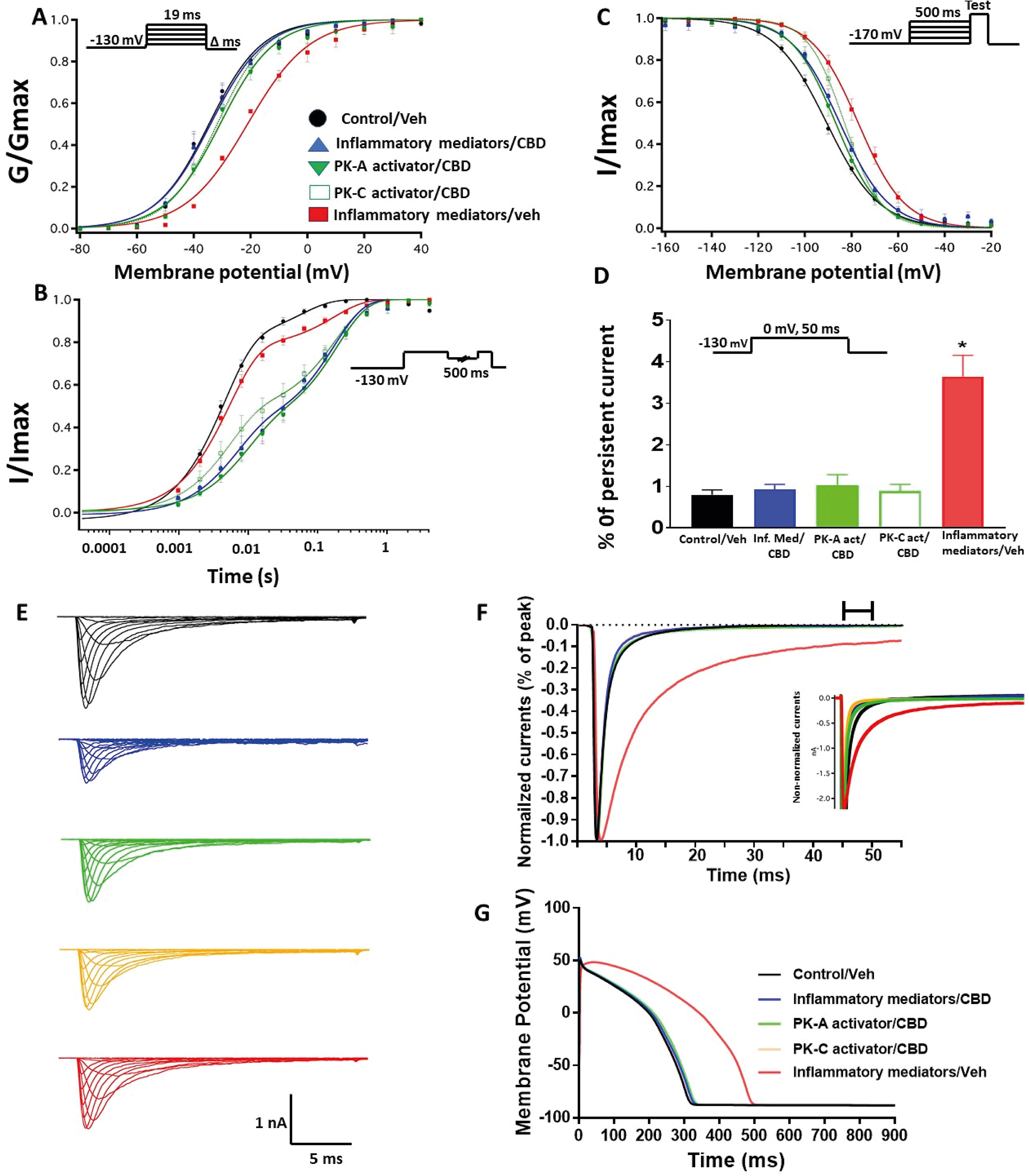
(A) Effect of CBD (5 μM, perfusion) on the conductance curve of Nav1.5 transfected cells incubated with inflammatory mediators (24 hours) or PK-A activator (CPT-cAMP; 1 μM, for 20 minutes) or PK-C activator (PMA; 10 nM, for 20 minutes) with the insert showing the protocol (n=5, each). (B) Effect of CBD (5 μM, perfusion) on SSFI of Nav1.5 transfected cells incubated with inflammatory mediators (24 hours) or PK-A activator (CPT-cAMP; 1 μM, for 20 minutes) or PK-C activator (PMA; 10 nM, for 20 minutes) with the insert showing the protocol (n=5, each). (C) Effect of CBD (5 μM, perfusion) on recovery from fast inactivation of Nav1.5 transfected cells incubated with inflammatory mediators (24 hours) or PK-A activator (CPT-cAMP; 1 μM, for 20 minutes) or PK-C activator (PMA; 10 nM, for 20 minutes) with the insert showing the protocol (n=5, each). (D) Effect of CBD (5 μM, perfusion) on the percentage of persistent sodium currents of Nav1.5 transfected cells incubated with inflammatory mediators (24 hours) or PK-A activator (CPT-cAMP; 1 μM, for 20 minutes) or PK-C activator (PMA; 10 nM, for 20 minutes) with the insert showing the protocol (n=5, each). (E) Representative families of macroscopic currents. (F) Representative persistent currents across conditions. Currents were normalized to peak current amplitude. Bar above current traces indicates period during which persistent current was measured. Inset shows non-normalized currents. (G) Effect of CBD (5 μM, perfusion) on the *In silico* action potential duration of Nav1.5 transfected cells incubated in inflammatory mediators (24 hours) or PK-A activator (CPT-cAMP; 1 μM, for 20 minutes) or PK-C activator (PMA; 10 nM, for 20 minutes). \**P* < 0.05 versus corresponding “Control/Veh” values.

### E_2_ rescues the high glucose-induced alterations in Nav1.5 gating via PK-A and PK-C pathway

We further investigated whether E_2_ rescues the high-glucose induced changes in biophysical properties of Nav1.5 given that E_2_ previously was shown to affect Nav in addition its anti-inflammatory role (Iorga, Cunningham, Moazeni, Ruffenach, Umar & Eghbali, 2017; Wang, Garro & Kuehl-Kovarik, 2010). We first tested the effects of E_2_ (5 or 10 μM) under control conditions and found that E_2_ exerted no significant effects on Nav1.5 gating (Tables 1–4). In contrast, Figure 5 shows that perfusing E_2_ (5 or 10 μM) for at least 10 minutes into the bath solution (Möller & Netzer, 2006; Wang et al., 2013) abolished the shifts elicited by high glucose (100 mM, for 24 hours, including V_1/2_, z of activation, and the V_1/2_ of SSFI in a concentration-dependent manner (Fig. 5A, 5B and Table 1 and 2). On the other hand, we found that E_2_ (5 or10 μM) had no significant effect on 100 mM glucose-induced slight increase in the slow component of fast inactivation recovery (Fig. 5C and Table 3). However, E_2_ significantly reduced the 100 mM glucose-induced increase in INap in a concentration-dependent manner (Fig. 5D and Table 4). E_2_ reduction of the glucose-exacerbated INap is consistent with previous reports of similar effects in neuronal sodium channels (Wang, Garro & Kuehl-Kovarik, 2010). Figure 5G shows AP modeling and suggests that E_2_, in a concentration-dependent manner, rescues the prolonged *in silico* APD caused by 100 mM glucose.

**Figure 5.**
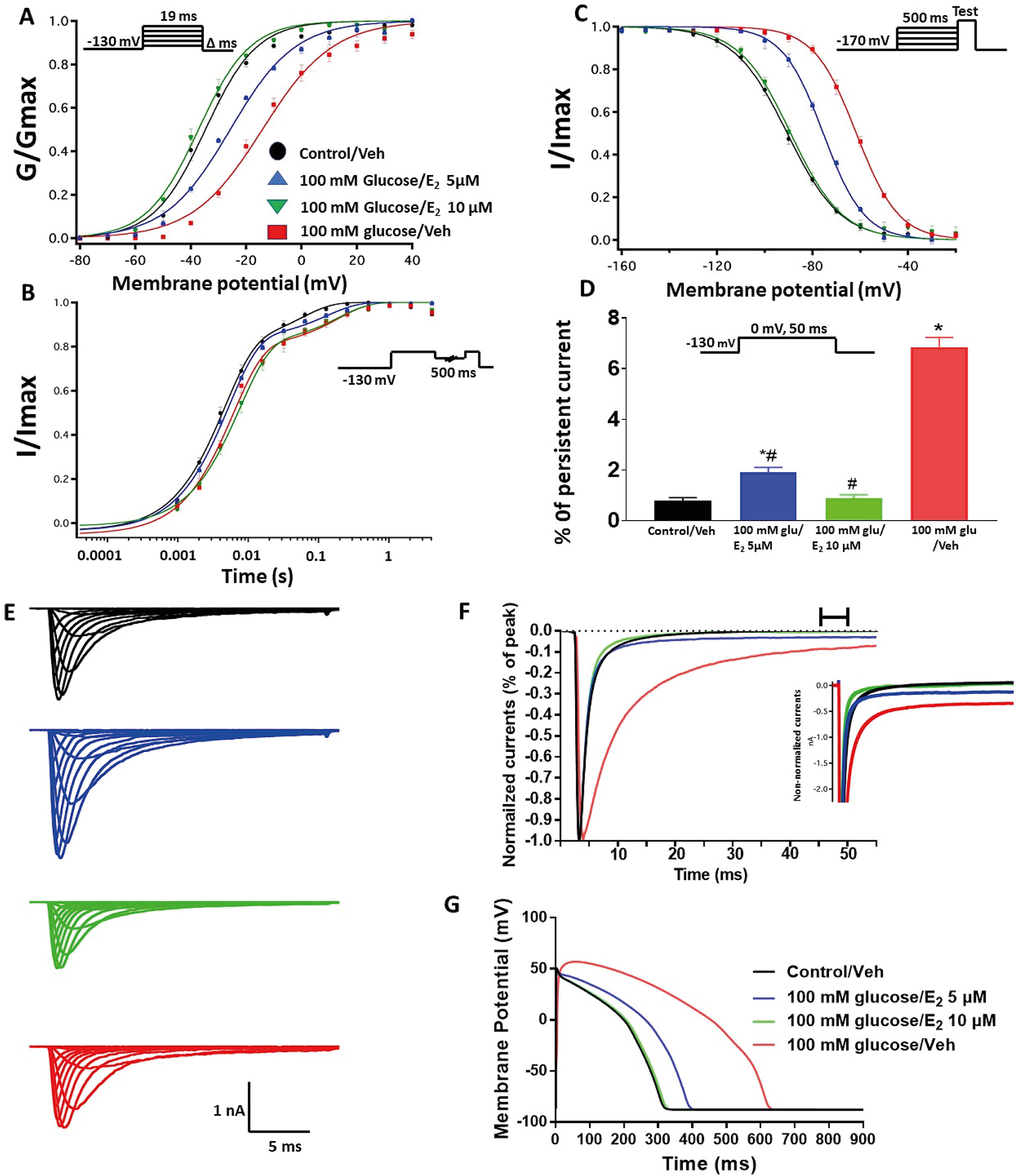
(A) Effect of E_2_ (5 or 10 μM) on conductance curve of Nav1.5 transfected cells incubated in 100 mM glucose (for 24 hours) with the insert showing the protocol (n=5, each). (B) Effect of E_2_ (5 or 10 μM) on SSFI of Nav1.5 transfected cells in 100 mM glucose (for 24 hours) with the insert showing the protocol (n=5, each). (C) Effect of E_2_ (5 or 10 μM) on recovery from fast inactivation of Nav1.5 transfected cells in 100 mM glucose (for 24 hours) with the insert showing the protocol (n=5, each). (D) Effect of E_2_ (5 or 10 μM) on the percentage of persistent sodium currents of Nav1.5 transfected cells in 100 mM glucose (for 24 hours) with the insert showing the protocol (n=5, each). (E) Representative families of macroscopic currents. (F) Representative persistent currents across conditions. Currents were normalized to peak current amplitude. Bar above current traces indicates period during which persistent current was measured. Inset shows non-normalized currents. (G) Effect of E_2_ (5 or 10 μM) on the *In silico* action potential duration of Nav1.5 transfected cells incubated in 100 mM glucose (for 24 hours). \**P* < 0.05 versus corresponding “Control/Veh” values. ^#^*P* < 0.05 versus corresponding “100 mM glucose/Veh” values.

We tested whether E_2_ (5 or 10 μM) rescues the effects of inflammatory mediators, PK-C activator (PMA), or PK-A activator (CPT-cAMP) on the gating properties of Nav1.5. Figure 6 shows that concurrent addition of E_2_ abolished the effects of inflammatory mediators on activation and SSFI in a concentration-dependent manner (Fig. 6A, 6B and Table 1 and 2). Similiarly, E_2_ concentration-dependently rescued PMA or CPT-cAMP-elicited effects on activation and SSFI (Fig. 6A, 6B and Table 1 and 2). Although E_2_ (5 or10 μM) had no significant effect on the slight increase in the slow component of fast inactivation recovery caused by inflammatory mediators, PMA, or CPT-cAMP (Fig. 6C and Table 3), E_2_ significantly reduced the increase in INap in a concentration-dependent manner (Fig. 6D and Table 4; representative currents shown in Figure 6E and 6F). In addition, E_2_ concentration-dependently rescues the prolonged *in silico* APD caused by inflammatory mediators or activators of PK-A or PK-C-induced to nearly that of the control condition (Fig. 6G).

**Figure 6.**
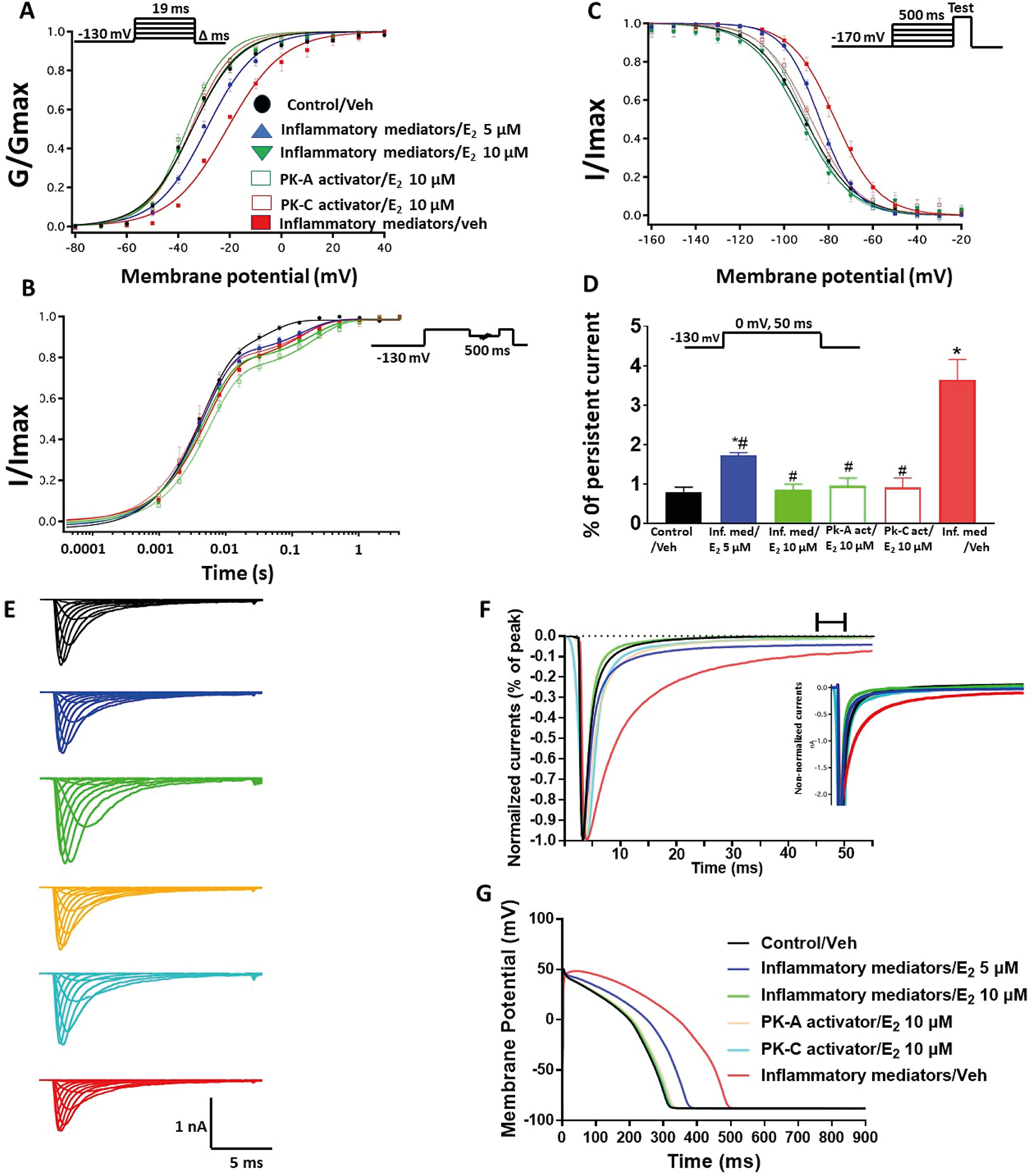
(A) Effect of E_2_ (5 or 10 μM) on conductance curve of Nav1.5 transfected cells incubated in inflammatory mediators (for 24 hours), PK-A activator (CPT-cAMP; 1 μM for 20 minutes) or PK-C activator (PMA; 10 nM, for 20 minutes) with the insert showing the protocol (n=5, each). (B) Effect of E_2_ (5 or 10 μM) on SSFI of Nav1.5 transfected cells incubated in inflammatory mediators (for 24 hours), PK-A activator (CPT-cAMP; 1 μM for 20 minutes) or PK-C activator (PMA; 10 nM, for 20 minutes) with the insert showing the protocol (n=5, each). (C) Effect of E_2_ (5 or 10 μM) on recovery from fast inactivation of Nav1.5 transfected cells incubated in inflammatory mediators (for 24 hours), PK-A activator (CPT-cAMP; 1 μM for 20 minutes) or PK-C activator (PMA; 10 nM, for 20 minutes) with the insert showing the protocol (n=5, each). (D) Effect of E_2_ (5 or 10 μM) on the percentage of persistent sodium currents of Nav1.5 transfected cells incubated in inflammatory mediators (for 24 hours), PK-A activator (CPT-cAMP; 1 μM for 20 minutes) or PK-C activator (PMA; 10 nM, for 20 minutes) with the insert showing the protocol (n=5, each). (E) Representative families of macroscopic currents. (F) Representative persistent currents across conditions. Currents were normalized to peak current amplitude. Bar above current traces indicates period during which persistent current was measured. Inset shows non-normalized currents. (G) Effect of E_2_ (5 or 10 μM) on the *In silico* action potential duration of Nav1.5 transfected cells incubated in inflammatory mediators (for 24 hours), PK-A activator (CPT-cAMP; 1 μM for 20 minutes) or PK-C activator (PMA; 10 nM, for 20 minutes). \**P* < 0.05 versus corresponding “Control/Veh” values. ^#^*P* < 0.05 versus corresponding “inflammatory mediators/Veh” values.

## Discussion

We recently showed that CBD confers protection on Nav1.5 against the high glucose-elicited hyperexictability and cytotoxicity (Fouda, Ghovanloo & Ruben, 2020). Here, we address, for the first time, the inflammation/PK-A and PK-C signaling pathway to mediate high glucose-induced cardiac anaomalies (Fig. 7). Our results suggest that CBD and E_2_ may exert their cardiprotective effects against high glucose, at least partly, through this signaling pathway. Our conclusions are based on the following main observations: (i) Similar to high glucose, inflammatory mediators elicited right shifts in the voltage-dependence of activation and inactivation, and exacerbated persistent currents. Increased persistent currents prolong the simulated action potential duration. (ii) Activators of PK-A and PK-C reproduced the high glucose- and inflammation-induced changes in Nav1.5 gating. (iii) Inhibitors of PK-A and PK-C reduced, to a great extent, the high glucose- and inflammation-induced changes in Nav1.5 gating. (iv) CBD or E_2_ rescued the effects of high glucose, inflammatory mediators, or PK-A or PK-C activators. Our results suggest a role for Nav1.5 in high glucose induced hyperexcitability, via inflammation and subsequent activation of PK-A and PK-C, which could lead to LQT3-type arrhythmia (Fig. 7). In addition, our findings suggest possible therapeutic effects for CBD in high glucose-provoked cardiac dysfunction in diabetic patients, especially those post-menopause. Diabetes-induced QT prolongation predisposes to malignant ventricular arrhythmias (Ukpabi & Onwubere, 2017). Moreover, LQT in diabetic patients make them three times more vulnerable to the risk of cardiac arrest (Whitsel et al., 2005). Nav1.5 gain-of-function plays a crucial role in the development of LQT (Shimizu & Antzelevitch, 1999). With that in mind, we found that inflammatory mediators replicated the high glucose-induced changes in Nav1.5 gating similar to those correlated with LQT3 in diabetic rats (Yu et al., 2018). This finding is consistent with other reports showing that hyperglycemia/high glucose is proinflammatory and that inflammation is a crucial player in the pathogenesis of cardiovascular anamolies (Fouda, Leffler & Abdel-Rahman, 2020; Tsalamandris et al., 2019). Accumulating evidence shows that inflammation is a potential cause for developing LQT through direct effects on myocardial electric properties, including its effect on Nav, and indirect autonomic cardiac regulations (Lazzerini, Capecchi & Laghi-Pasini, 2015). Inflammation alters the electrophysiological properties of cardiomyocytes Nav with an increase in INap leading to prolongation of APD, similar to our findings (Fig. 1) (Shryock, Song, Rajamani, Antzelevitch & Belardinelli, 2013; Ward, Bazzazi, Clark, Nygren & Giles, 2006). Taken together, these findings support our hypothesis that high glucose, at least partly through induction of inflammation, alters Nav1.5 gating and leads to LQT arrhythmia (Fig.7).

**Figure 7.**
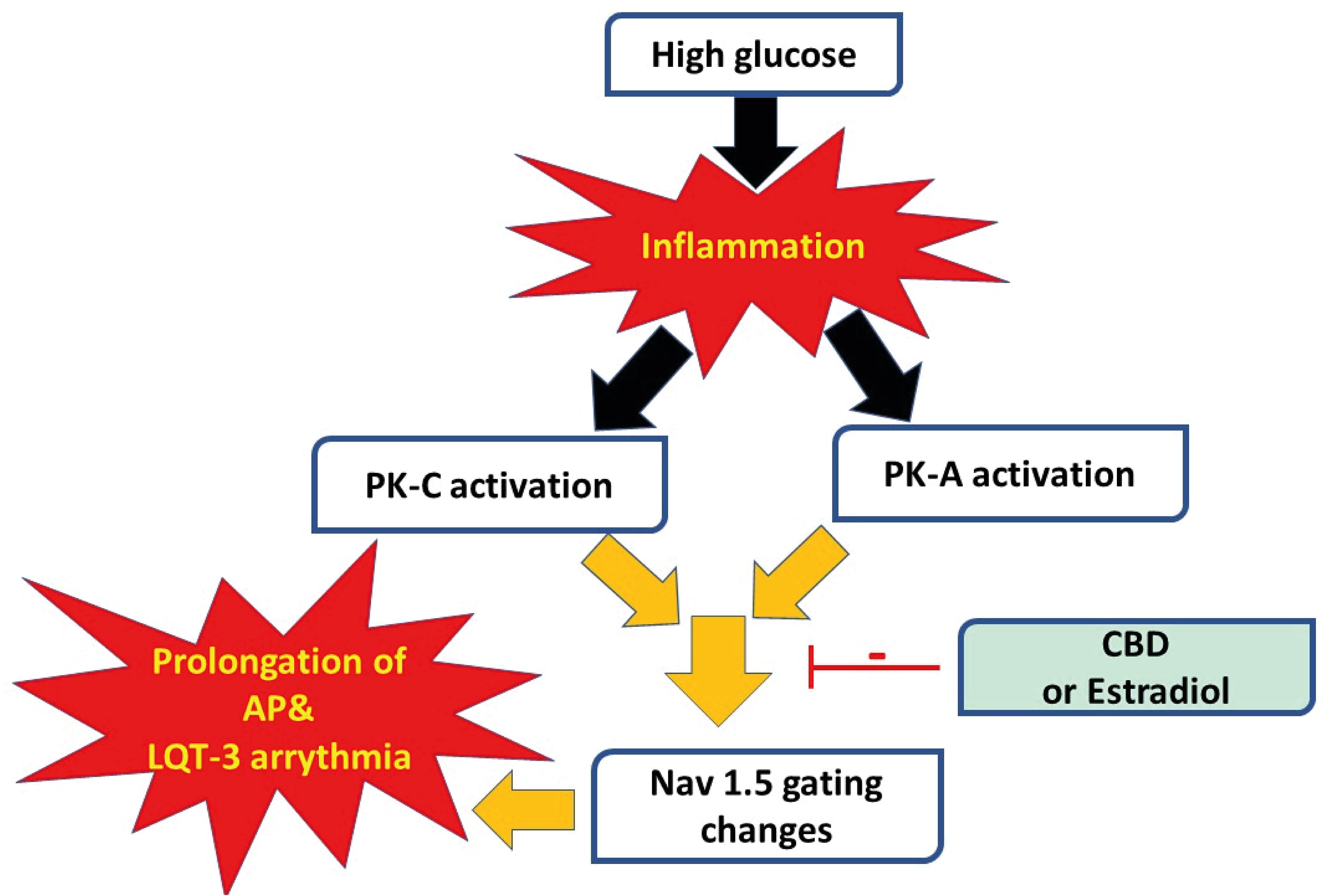
A schematic of possible cellular pathway involved in the protective effect of CBD, E_2_ against high glucose induced inflammation and activation of PK-A and PK-C via affecting cardiac voltage-gated sodium channels (Nav1.5).

The activation of PK-A and PK-C and subsequent protein phosphorylation is among the key signaling pathways associated with inflammation (Karin, 2005) and hyperglycemia, resulting in many devastating diabetes-induced cardiac complications (Bockus & Humphries, 2015; Koya & King, 1998). Our data suggest that activation of PK-A or PK-C replicated high glucose- and inflammation-induced gating changes in Nav1.5 gating, whereas inhibition of PK-A or PK-C abolished those changes (Figs. 2 and 3). This finding suggests that PK-A and PK-C may be downstream effectors of inflammation in high glucose-induced cardiac complications (Fig. 7). PK-A phosphorlylates S525 and S528, while PK-C phosphorylates S1503 in human Nav1.5 (Iqbal & Lemmens-Gruber, 2019). There are conflicting reports regarding the effects of PK-A and PK-C activation on the voltage-dependence and kinetics of Nav1.5 gating. These differences could be attributed to different voltage protocols, different holding potentials, different concentrations or type of PK-activators, or different cell lines used in the various studies (Aromolaran, Chahine & Boutjdir, 2018; Iqbal & Lemmens-Gruber, 2019). Despite this discrepancy, both PK-A or PK-C destabilize Nav fast inactivation and hence increase INap, which is strongly correlated to prolonged APD as shown in our findings (Fig. 7) (Astman, Gutnick & Fleidervish, 1998; Franceschetti, Taverna, Sancini, Panzica, Lombardi & Avanzini, 2000; Tateyama, Rivolta, Clancy & Kass, 2003).

Our results with PK-A and PK-C modulators prompted us to test whether CBD affects the biophysical properties of Nav1.5 through this pathway. We investigated the possible protective effect of CBD against the deletrious effects of high glucose through this signaling pathway because CBD protects against high glucose-induced gating changes in Nav1.5 (Fouda, Ghovanloo & Ruben, 2020). In addition, CBD attenuates the diabetes-induced inflammation and subsequent cardiac fibrosis through inhibition of phosphorylation enzymes (such as MAPKs) (Rajesh et al., 2010). Our results suggest that CBD alleviates the inflammation/activation of PK-A or PK-C induced biophyscial changes (Fig. 4). Our findings are consistent with the anti-inflammatory, anti-oxidant, and anti-tumor effects of CBD via inhibition of PK-A and PK-C signaling (Seltzer, Watters & MacKenzie, 2020). The incomplete protective effects of PK-A and PK-C inhibitors compared to the CBD effect against the inflammation-induced gating changes in Nav1.5 could be attributed to the combined CBD direct inhibtory effect on Nav1.5 and its indirect inhibitory actions on both PK-A and PK-C (Figs. 3, 4 and 7).

Interestingly, E_2_ directly affects Nav and exerts anti-inflammatory effects (Iorga, Cunningham, Moazeni, Ruffenach, Umar & Eghbali, 2017; Wang, Garro & Kuehl-Kovarik, 2010). We found that E_2_, similar to CBD, rescues the effects of high-glucose, inflammation, and activation of PK-A or PK-C (Figs. 5–7). Our results are consistent with other reports showing the cardioprotective effects of E_2_ by increasing angiogenesis, vasodilation, and decreasing oxidative stress and fibrosis (Iorga, Cunningham, Moazeni, Ruffenach, Umar & Eghbali, 2017). Although the role of E_2_ in arrhythmias is controversial, many studies support the anti-arrythmic effects of E_2_ because of its effects on the expression and function of cardiac ion channels (Iorga, Cunningham, Moazeni, Ruffenach, Umar & Eghbali, 2017; Odening & Koren, 2014). Notably, E_2_ stabilizes Nav fast inactivation and reduces INap, similar to CBD (Wang, Garro & Kuehl-Kovarik, 2010). Further, E_2_ reduces the oxidative stress and the inflammatory reponses by inhibiting PK-A and PK-C-mediated signaling pathways (Mize, Shapiro & Dorsa, 2003; Viviani, Corsini, Binaglia, Lucchi, Galli & Marinovich, 2002).

In conclusion, our results suggest that inflammation and the subsequent activation of PK-A and PK-C correlate with the high glucose-induced electrophysiological changes in Nav1.5 gating (Fig. 7). *In silico*, these gating changes result in prolongation of simulated action potentials leading to LQT3 arrhythmia, which is a clinical complication of diabetes (Grisanti, 2018). CBD and E_2_, through inhibition of this signaling pathway, ameliorate the effects of high glucose and the resultant clinical condition. In light of the debate about the risks associated with hormonal replacement therapy (Climént-Palmer & Spiegelhalter, 2019), CBD may provide an alternate therapeutic approach, especially in diabetic post-menopausal populations due to their decreased levels of cardioprotective E_2_ (Xu, Lin, Wang, Xiong & Zhu, 2014).

## Acknowledgments

This work was supported by MITACS Elevate grant in partnership with Akseera Pharma, Inc. (IT14450) to MAF.

The authors thank Dr. Mohammad-Reza Ghovanloo for his help in the action potential modelling, and Dr. Dana Page for her thoughtful contributions to the manuscript.

## Author and contributors

MAF collected, assembled, analyzed, interpreted the data and wrote the first draft of the manuscript. PCR conceived the experiments and revised the manuscript critically for important intellectual content.

## Declaration of transparency and scientific rigour

This Declaration acknowledges that this paper adheres to the principles for transparent reporting and scientific rigour of preclinical research recommended by funding agencies, publishers and other organisations engaged with supporting research.

